# Modeling within-host and aerosol dynamics of SARS-CoV-2: the relationship with infectiousness

**DOI:** 10.1101/2022.03.08.483569

**Authors:** Nora Heitzman-Breen, Stanca M. Ciupe

**Affiliations:** Department of Mathematics, Virginia Polytechnic Institute and State University, Blacksburg, VA, USA

## Abstract

The relationship between transmission of severe acute respiratory syndrome coronavirus 2 (SARS-CoV-2) and the amount of virus present in the proximity of a susceptible host is not understood. Here, we developed a within-host and aerosol mathematical model and used it to determine the relationship between viral kinetics in the upper respiratory track, viral kinetics in the aerosols, and new transmissions in golden hamsters challenged with SARS-CoV-2. We determined that infectious virus shedding early in infection correlates with transmission events, shedding of infectious virus diminishes late in the infection, and high viral RNA levels late in the infection is a poor indicator of transmission. We further showed that viral infectiousness increases in a density dependent manner with viral RNA and that their relative ratio is time-dependent. Such information is useful for designing interventions.

**Author summary:** Quantifying the relationship between SARS-CoV-2 dynamics in upper respiratory tract and in aerosols is key to understanding SARS-CoV-2 transmission and evaluating intervention strategies. Of particular interest is the link between the viral RNA measured by PCR and a subject’s infectiousness. Here, we developed a mechanistic model of viral transmission in golden hamsters and used data in upper respiratory tract and aerosols to evaluate key within-host and environment based viral parameters. The significance of our research is in identifying the timing and duration of viral shedding, how long it stays infectious, and the link between infectious virus and total viral RNA. Such knowledge enhances our understanding of the SARS-CoV-2 transmission window.

## 1 Introduction

The transmission of the severe acute respiratory syndrome coronavirus 2 (SARS-CoV-2), the agent that causes coronavirus disease 2019 (COVID-19), is dependent on the amount of infectious particles present in the environment surrounding the susceptible host, and/or on the proximity between the susceptible and infectious host. Experimental studies that use real-time reverse transcription-polymerase chain reaction (PCR) assays have reported the presence of SARS-CoV-2 RNA in contaminated environmental surfaces [3, 12, 19, 22] and in aerosols [17, 20, 21, 24]. Moreover, *in vivo* cell culture assays, have shown that particle released into the environment are replication-competent [33] and they can stay infectious in the aerosols for up to three hours [28].

Once an infection is established, the dynamics of SARS-CoV-2 in the upper respiratory tract (URT) is dictated by the interplay between the virus fitness and host immune responses. The total RNA to infectious virus ratio changes over the course of the infection, varying between 10^3^ : 1 and 10^6^ : 1 RNA to plaque forming units (PFU) [16, 30]. Determining the within-host mechanistic interactions responsible for the temporal changes in infectious to non-infectious viral dynamics is important for guiding interventions.

Over the last two years, within-host mathematical models developed for influenza and other respiratory infections have been modified for SARS-CoV-2 infections [10, 11, 13, 15, 23, 31]. These models divided total viral titers into infectious and non-infectious particles [7, 27], fitted their sum to total RNA values measured by PCR (used as a proxy for total virus load) and used the results to determine the mechanisms of viral expansion and loss. The models were subsequently used to provide insights into the relative fitness of variants, types of drug interventions [13, 15, 23], the relationship between individual infection and population transmission, and the effect of this relationship on testing and vaccine strategies [5, 6]. In most studies, the total RNA to infectious virus ratio in URT is assumed constant over time. However, Goyal *et al*. [8] and Ke *et al*. [14] proposed a nonlinear correspondence between infectious virus and total RNA, and found that a density dependent function best describes their relationship [14].

While there is a reasonable understanding of the mechanistic dynamics modulating SARS CoV-2 infection in the URT, the temporal shedding into the environment of viral RNA and infectious virus has not been explored. Here, we expand a within-host mathematical model to include the dynamics of viral RNA and infectious virus titers in both URT and aerosols. We validate the models against two URT and one aerosol inoculation study in golden hamsters [9, 25] and use the models to determine the relationship between infectious virus an total RNA in both environments. Lastly, we investigate the link between the infectious virus shed in the environment and the probability of a nearby host getting infected. The results can guide interventions.

## 2 Material and methods

### Experimental data

We use previously published temporal SARS-CoV-2 RNA and infectious virus titer data from two inoculation studies in golden Syrian hamsters:

- *Sia et al. study [25]* : six donor hamsters (all male) were inoculated intranasally with 8 × 10^4^ tissue culture ineffective dose (TCID50) of SARS-CoV-2. At 24 hours after inoculation, each donor was transferred to a new cage and co-housed with one naive hamster. Viral RNA and infectious virus titers were collected every other day (in RNA/ml and TCID50/ml) for the first 14 days in both donors and contacts and their weight changes were monitored daily. We will refer to these two groups as donors and contacts.
- *Hawks et al. study [9]:* eight hamsters (4 males and 4 females) were inoculated intranasally with 10^5^ PFU (1.4 × 10^5^ TCID50) of SARS-CoV-2. Viral RNA and infectious virus titers were collected daily from nasal washes (in RNA/wash and PFU/wash) and from the exhaled breath (aerosols) (in RNA/hour and PFU/hour) for the first five days and then again at day 10. Since the nasal wash and air samples were collected in 100*µl* and 400*µl*, we rescaled the RNA/wash data to RNA/ml by multiplying it with 10 and 2.5, respectively. Moreover, since 1 TCID50= 0.7 PFU, we rescaled the PFU/wash infectious virus titers in the nasal washes and in the air to TCID50/ml by multiplying them with 10*/*0.7 and 2.5*/*0.7, respectively. Hamsters were weighed daily and the study was terminated when clinical signs of illness were observed. We will refer to these two groups as males and females.

### Within-host and aerosol model

We model the interaction between target epithelial cells *T* , exposed epithelial cells *E*, infected epithelial cells *I*, infectious virus in upper respiratory tract *V*_*u*_, infectious virus in the air *V*_*a*_, total viral RNA in upper respiratory tract *R*_*u*_, and total viral RNA in the air *R*_*a*_ (see **Fig. 1** for a description). We assume that target cells get infected at rate *β* and become productively infected at rate *k*. Productively infected cells produce infectious virus at rate *p* and die at rate *δ*. Infectious virus particles in the upper respiratory tract are removed at rate *d* + *c*, where *d* is degradation rate and *c* is the immune induced clearance rate. Infectious virus is shed into the air at rate *ϕ*_1_, where it loses infectiousness at rate *d* + *d*_1_, where *d* is the degradation rate (as before) and *d*_1_ accounts for enhanced inactivation due to the elements. The equations describing these interactions are

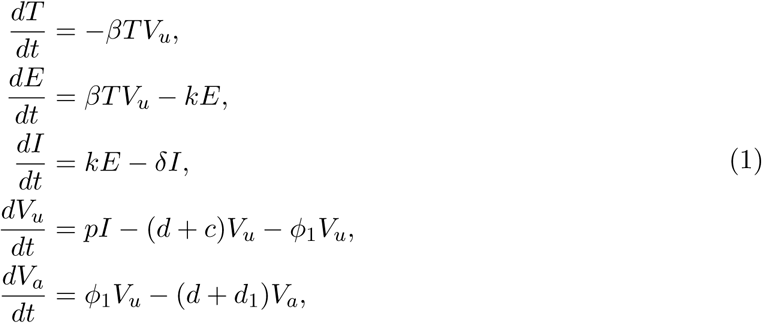

with initial conditions *T*(0) = *T*_0_, *E*(0) = 0, *I*(0) = 0, *V*_*u*_(0) = *V*_0_, *V*_*a*_(0) = 0.

**Figure 1:**
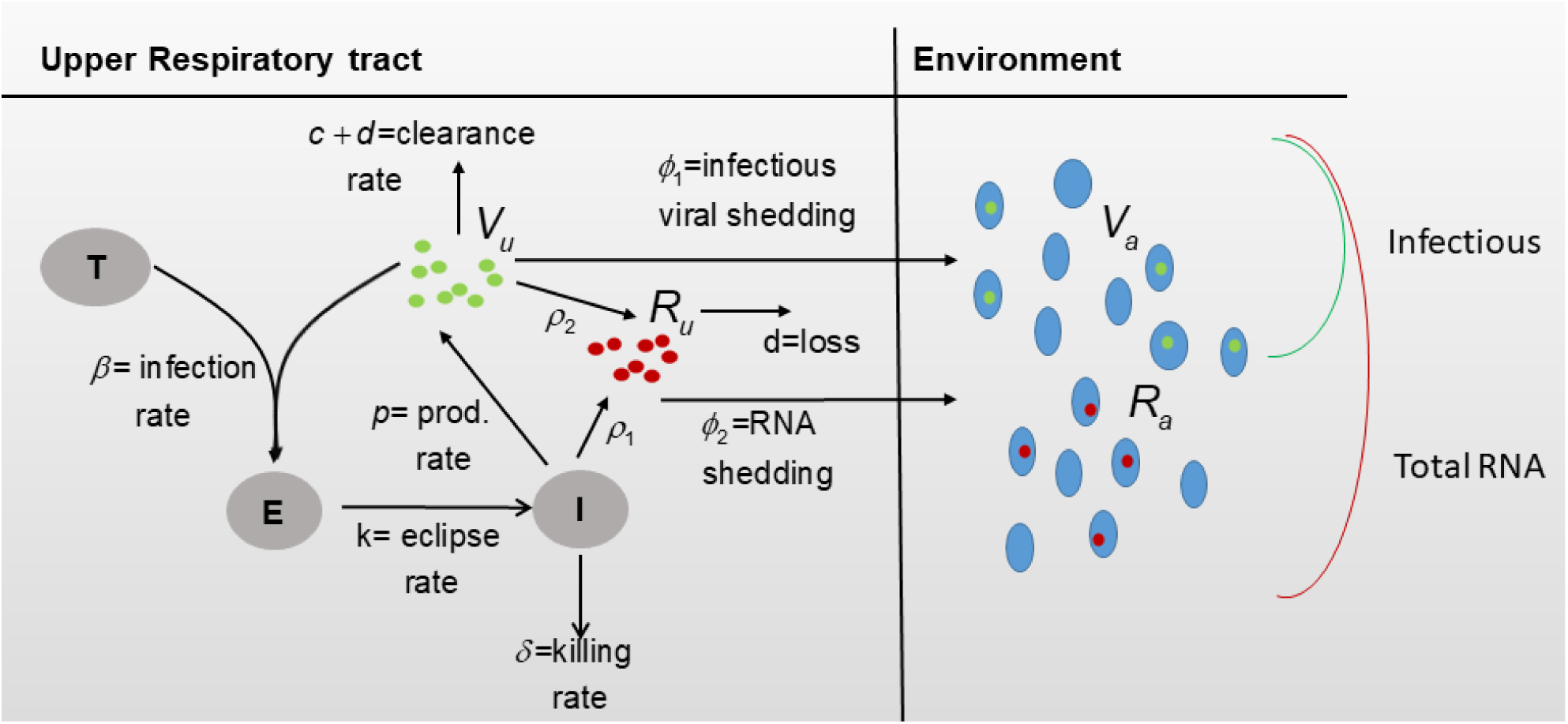
Diagram for model Eqs. (3) and (4).

Moreover, we model viral RNA in the upper respiratory tract *R*_*u*_ and in the air *R*_*a*_ as follows. Since the measured viral RNA may not be a good representation of actual virions (infectious or noninfectious) but also of naked viral RNA from virus neutralization or infected cells death, we assume viral RNA is produced at rate *ρ*_1_ per infected cell, and rate *ρ*_2_ per infectious virus. Rate *ρ*_1_ incorporates both newly synthesized genomic RNA that can be used for replication or transcription and naked RNA released from dead infected cells. Rate *ρ*_2_ represents neutralized infectious virus. The equations describing the viral RNA dynamics over time are given by

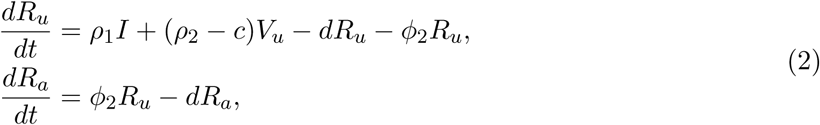

with initial conditions *R*_*u*_(0) = *R*_*a*_(0) = 0.

### Parameter values

We assume an initial target cell population in the upper respiratory tract *T*(0) = 10^7^ epithelial cells/ml, as in influenza [26], and no exposed and infected cells *E*(0) = *I*(0) = 0 epithelial cells/ml. The initial infectious virus is given by the inoculum titer, *V*_*u*_(0) = 8 × 10^4^ TCID50 for donors and contacts and *V*_*u*_(0) = 1.4 × 10^5^ TCID50 for males and females. No viral RNA is present in URT at the time of inoculation, *V*_*a*_(0) = 0 TCID50, and neither infectious virus nor viral RNA are present in the air at the time of inoculation, *R*_*u*_(0) = *R*_*a*_(0) = 0 TCID50. We assume that the infectious virus removal rate is *c* + *d* = 10 per day [13], the eclipse rate is *k* = 4 per day [14], the RNA degradation rate is *d* = 1 per day, and the RNA release due to neutralization is *ρ*_2_ = *c* = 9 per day. Lastly, since RNA and infectious virus from exhaled breath are only collected in Hawks *et al*. [9], we assume *ϕ*_1_ = *ϕ*_2_ = 0 in models (1) and (2) when applied to donors and contacts and {*ϕ*_1_, *ϕ*_2_} ≠ 0 when applied to males and females.

### Incorporating weight variability into the model

The weights of males and females vary from 56 to 76.1 grams (with an average of 70 grams) at inoculation. After challenge, each subject was weighed, as a way to monitor them for clinical signs of illness, and daily percent weight changes were reported as changes from the baseline weight *w*_0_ = 1, 0 ≤ *w*_*i*_ ≤ 1 for days *i* ∈ *S* = {1, 2, 3, 4, 5, 10} post infection. To account for all intermediate time points, we fitted a 4-degree polynomial to the weight data (smallest degree polynomial that gave a residual sum of squares < 10^−3^), and the resulting percent weight functions are shown in **Fig. 2A**.

**Figure 2:**
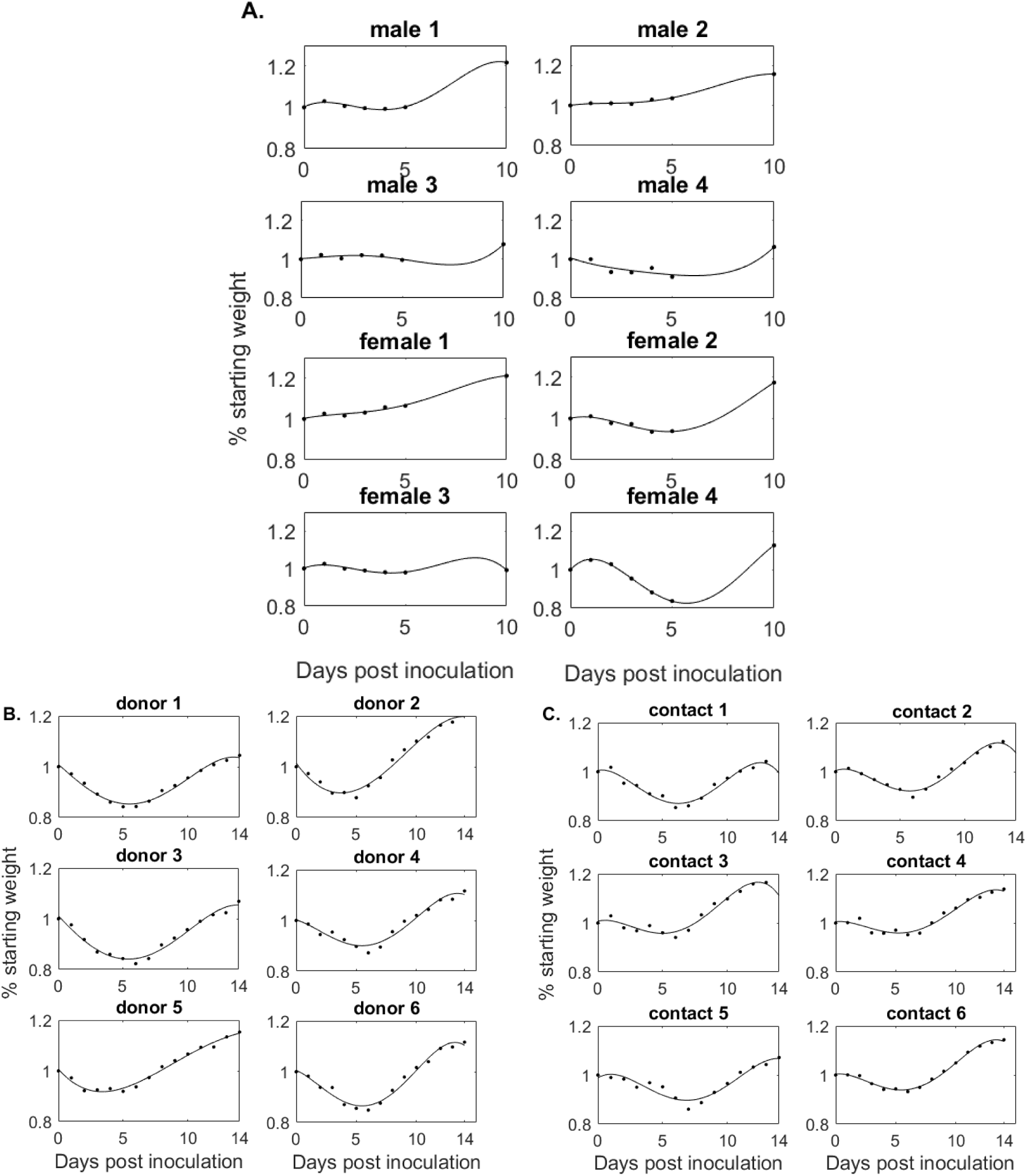
% Weight change from baseline over time (black lines) versus weight data (diamonds) in (A) males and females, (B) donors, and (C) contacts.

The initial weights of donors and contacts are unknown. After inoculation, each subject was weighed and percent weight changes were reported as changes from the baseline weight *w*_0_ = 1, 0 ≤ *w*_*i*_ ≤ 1 for days *i* ∈ *S* = {2, 4, 6, 8, 10, 12, 14} post infection. As before, we fitted a 4-degree polynomial to the weight data, and the resulting percent weight functions are shown in **Fig. 2B** and **Fig. 2C** for the donors and contacts, respectively.

To include the weight variability into the model, we assume that weight changes result in adjusted available epithelial cells. At day *t* target, exposed and infected cells become *T*_1_(*t*) = *T*(*t*) × *w*(*t*), *E*_1_(*t*) = *E*(*t*) × *w*(*t*), and *I*_1_(*t*) = *I*(*t*) × *w*(*t*). The models for the weight-dependent cell populations become

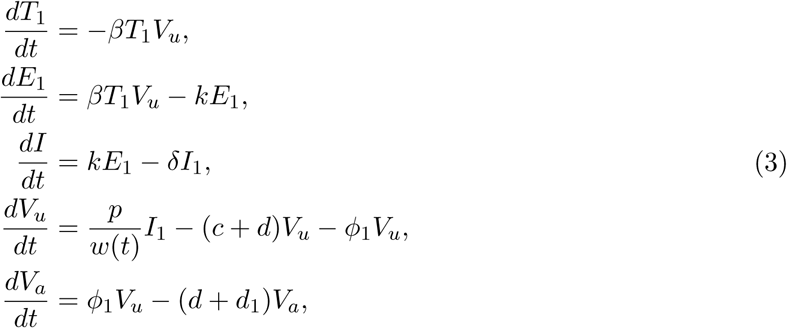

and

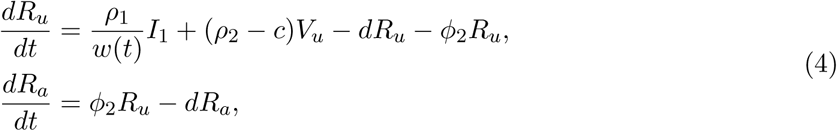

Moreover, the initial conditions become *T*_1_(0) = *ξT*_0_, *E*_1_(0) = 0, *I*_1_(0) = 0, *V*_*u*_(0) = *V*_0_, *V*_*a*_(0) = 0, *R*_*u*_(0) = *R*_*a*_(0) = 0 where *ξ* = {1.06, 1.09, 0.97, 0.81, 1.01, 0.92, 1.094, 1.04} is scaling from the average initial weight of 70 grams for males and females and *ξ* = 1 for donors and contacts [25].

### The basic reproduction number

The basic reproduction number (or basic reproductive ratio) is defined as the number of infected cells (or virus particles) that are produced by one infected cell (or virus particle) when the virus is introduced into a population of uninfected target cells *T*_1_(0). It is given by

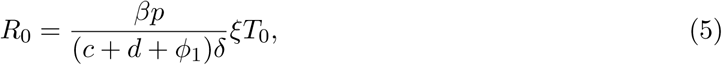

with *ϕ*_1_ = 0 for donors and contacts.

### Data fitting

Using models (3) and (4) we estimate parameters 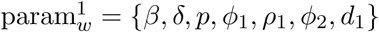, by minimizing the functional

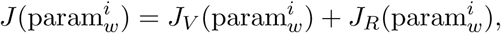

where

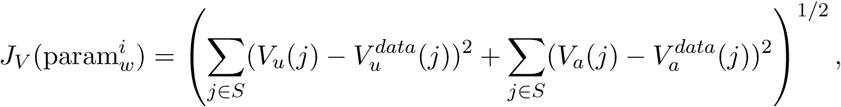

and

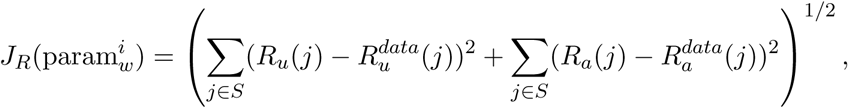

where *S* = {1, …, 5, 10} days post infection.

Using model (3) we estimate parameters 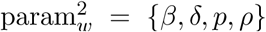 for donors and 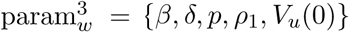 for contacts by minimizing the functional

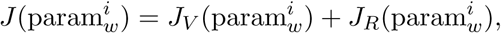

where

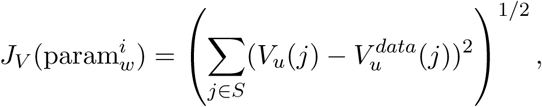

and

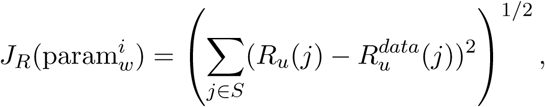

where *S* = {2, 4, 6, 8, 10, 12, 14} days post infection for donors and *S* = {1, 3, 5, 7, 9, 11, 13} days post infection for contacts. We only consider the first data point at or below limit of detection. We use the ’fminserach’ algorithm in matlab and the resulting estimates are given in **Tables 1** and **2**. The theoretical solutions of *V*_*u*_ and *R*_*u*_ versus male and female data are shown in the left panels of **Fig. 3** and versus donor and contact data are shown in **Fig. 4**.

**Table 1:** Individual estimates from simultaneously fitting *V*_*u*_, *V*_*a*_ and *R*_*u*_, *R*_*a*_ given by models Eqs. (3) and (4) to URT and aerosol infectious virus and RNA data in the males and female groups in Hawks *et al*. [9].

| Subject | $\beta \times 10^{-6}$ | $p$ | $\delta$ | $\phi_1 \times 10^{-4}$ | $\phi_2 \times 10^{-4}$ | $\rho_1$ | $d_1$ | ssq |
| --- | --- | --- | --- | --- | --- | --- | --- | --- |
| male 1 | 2.09 | 18.92 | 5.44 | 4.74 | 1.53 | 155.91 | 3.45 | 4.26 |
| male 2 | 6.46 | 6.41 | 4.44 | 3.29 | 1.14 | 106.12 | 1.58 | 3.94 |
| male 3 | 3.74 | 61.41 | 3.47 | 0.13 | 0.03 | 183.02 | 0.90 | 2.51 |
| male 4 | 14.2 | 2.48 | 1.21 | 6.33 | 0.58 | 66.69 | 3.94 | 4.54 |
| average | 6.62 | 22.30 | 3.64 | 3.62 | 0.82 | 127.93 | 2.47 |  |
| female 5 | 40.6 | 0.03 | 1.91 | 21.0 | 2.56 | 14.38 | 0.00 | 6.80 |
| female 6 | 19.2 | 0.23 | 5.41 | 14.0 | 1.26 | 167.32 | 1.71 | 5.52 |
| female 7 | 17.5 | 0.10 | 3.95 | 28.0 | 0.17 | 335.85 | 0.00 | 5.33 |
| female 8 | 25.0 | 0.06 | 2.50 | 13.0 | 0.09 | 341.59 | 0.94 | 4.10 |
| average | 25.6 | 0.10 | 3.44 | 19.0 | 1.02 | 214.79 | 0.66 |  |

**Table 2:** Individual estimates from simultaneously fitting *V*_*u*_ and *R*_*u*_ given by model Eq. (3) to URT infectious virus and RNA data in donors and contacts from Sia *et al*. [25].

| Subject | $\beta \times 10^{-6}$ | $p$ | $\delta$ | $\rho_2 \times 10^4$ | $V_0$ | ssq |
| --- | --- | --- | --- | --- | --- | --- |
| Donor 1 | 3.84 | 16.8 | 2.60 | 22.92 | - | 1.19 |
| Donor 2 | 4.42 | 19.7 | 2.54 | 7.14 | - | 1.64 |
| Donor 3 | 3.89 | 19.2 | 2.56 | 14.41 | - | 1.10 |
| Donor 4 | 5.79 | 13.2 | 1.82 | 2.68 | - | 2.16 |
| Donor 5 | 8.52 | 3.50 | 1.78 | 0.28 | - | 2.94 |
| Donor 6 | 5.76 | 21.0 | 1.98 | 1.16 | - | 3.01 |
| Donor Av. | 5.37 | 15.6 | 2.21 | 8.10 | - |  |
| Contact 1 | 21.5 | 0.96 | 2.12 | 5.45 | 171 | 0.84 |
| Contact 2 | 12.4 | 3.49 | 2.53 | 9.47 | 1.43 | 2.51 |
| Contact 3 | 40.5 | 0.72 | 1.81 | 7.05 | 283 | 0.99 |
| Contact 4 | 2.42 | 7.54 | 4.11 | 1.44 | 969 | 1.59 |
| Contact 5 | 46.8 | 0.55 | 2.02 | 0.81 | 55.8 | 1.82 |
| Contact 6 | 0.54 | 44.5 | 4.38 | 2.73 | 937 | 1.98 |
| Contact Av. | 20.7 | 9.62 | 2.82 | 4.49 | 402 |  |

**Figure 3:**
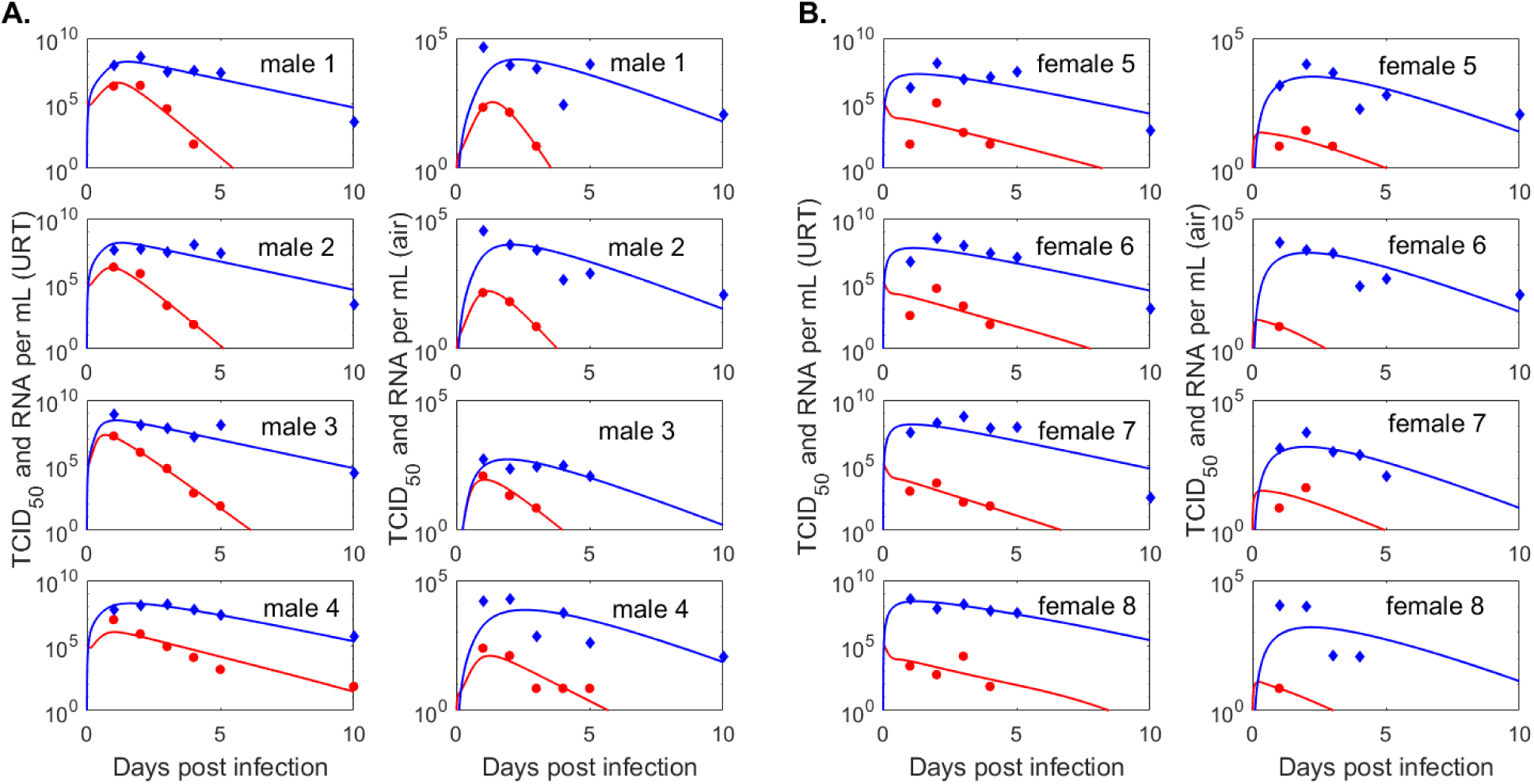
(Left panels) Dynamics of infectious virus *V*_*u*_ (red lines) and viral RNA *R*_*u*_ (blue lines) as given by model Eq. 3 versus infectious viral titers (red circles) and RNA (blue diamonds) in the upper respiratory tract of the (A.) males and (B.) females; (Right panels) Dynamics of infectious virus *V*_*a*_ (red lines) and RNA molecules *R*_*a*_ (blue lines) as given by model Eq. 4 versus infectious viral titers (red circles) and RNA (blue diamonds) in the exhaled breath of (A.) males and (B.) females. Model parameters are given in Table 1.

**Figure 4:**
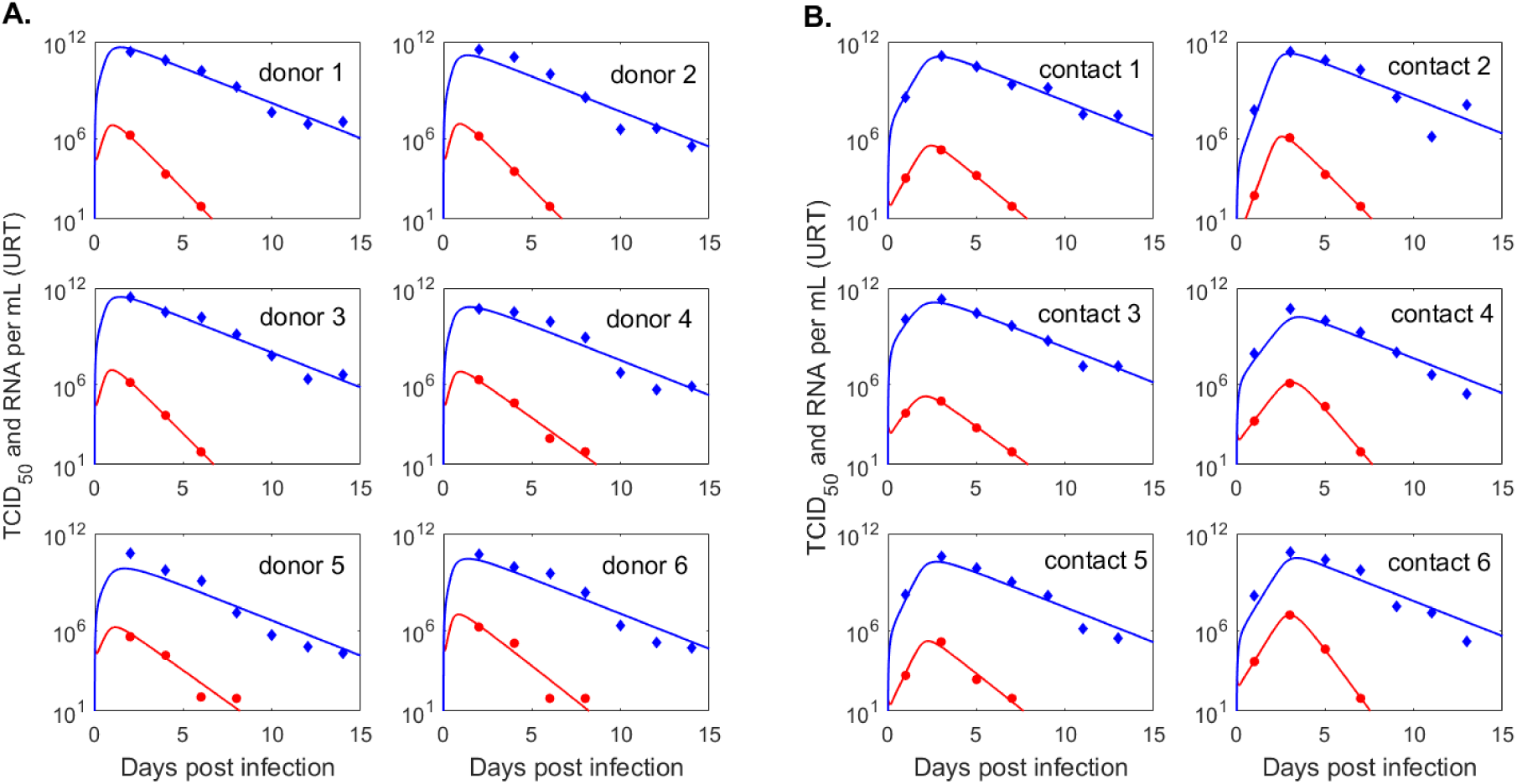
Dynamics of infectious virus *V*_*u*_ (red lines) and viral RNA *R*_*u*_ (blue lines) as given by model Eq. 3 versus infectious viral titers (red circles) and viral RNA (blue diamonds) in the upper respiratory tract of (A.) donors and (B.) contacts. Model parameters are given in Table 2.

### Modeling the relationship between infectious virus titer and total RNA

We assume that the log10 URT infectious virus titer, *ν* = log_10_ *V*_*u*_ (measured in log10 TCID50 per mL) can be modeled as a logistic growth function of the log10 total viral RNA, *ρ* = log_10_ *R*_*u*_ (measured in log10 RNA per mL), with per capita growth rate *r* and carrying capacity *L*

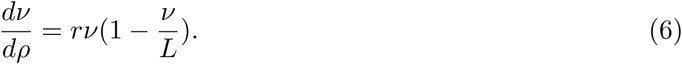

Given an initial infectious virus titer, *ν*(0) = *ν*_0_, we obtain the following solution for the log10 infectious virus as a function of total RNA (identical with the density dependent functions considered by Ke *et al*. [13] and Goyal *et al*. [8])

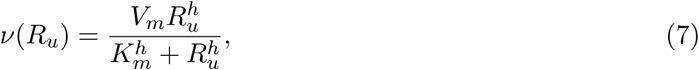

where 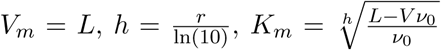. Here, *ν* is in units log10 TCID50 per mL and *R*_*u*_ is in units RNA per mL. We estimate the distribution of parameters 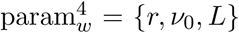 by fitting equation (6) to the donors, contacts and males (*ν, R*_*u*_) data. We excluded the female group due to limited infectious virus titers above the limit of detection. All data at and below limit of detection is treated as censored during data fitting. We maximize the likelihood,

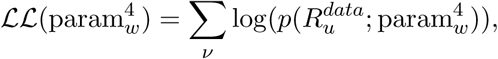

using a stochastic approximation expectation-maximization (SAEM) algorithm with constant residual errors implemented in Monolix 2019r2 [1]. Here, *p* represents the probability distribution governing the observation of infectious virus, 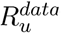. We assume a nonlinear mixed effects model that applies log-normal distributions for *r* and *ν*_0_ and logit-normal distribution with domain (0, 10] for *L*, to ensure biologically reasonable values for *V*_*m*_. Additionally, for the male group, we further assume a logit-normal distribution for *ν*_0_ with domain (0, 1 × 10^−3^] (to further ensure biological results for *K*_*m*_). This approach maximizes the likelihood that the estimated parameter distributions represent the true distributions for the given data. We establish a goodness of fit by calculating the Akaike Information criterion estimates

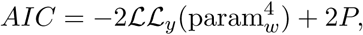

where 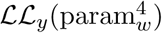 is the log-likelihood estimate for population parameters 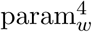 obtained using the importance sampling Monte Carlo method in Monolix 2019r2 [1] and *P* = 7 is the number of parameters estimated (including the mean and shape for 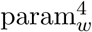 and constant error parameter). The resulting population parameter distributions and corresponding AICs are given **Table 3** and the population fits for log10 infectious virus *ν* = log_10_ *V*_*u*_ versus total viral RNA *R*_*u*_ for each group are shown in **Fig. 7A-C**.

**Table 3:** Expected population mean parameters and associated AIC values from fitting Eq. (6) to infectious versus RNA data from donors, contacts and males.

| | $V_m$ | $h$ | $K_M$ | AIC |
| --- | --- | --- | --- | --- |
| Donors | 8.7039 | 0.61 | $5.04 \times 10^{10}$ | 69.43 |
| Contacts | 8.9879 | 0.59 | $6.00 \times 10^{10}$ | 104.98 |
| Males | 8.2662 | 0.60 | $6.53 \times 10^7$ | 89.03 |

## 3 Results

### Viral kinetics in the upper respiratory tract

To study the kinetics of infectious virus titers and total viral RNA in the upper respiratory tract we used a dynamical within-host target cell limitation model developed for other acute respiratory infections (**Eq. 3**), which was normalized to include the temporal change in the subject’s weight (see **Materials and Methods** for a full description). We estimated unknown biological parameters by fitting the model to infectious viral titer and total RNA data of four groups (males, females, donors, contacts) from two inoculation studies [9,25] (see **Materials and Methods**). The resulting dynamics are in good agreement with the data kinetics in all four groups, with some inter group differences in the predicted outcomes. Infectious virus titers in the male and donors groups, who were challenged with high viral dose, expand to reach a peak 12 − 24 hours after inoculation and decline below limit of detection by 5-8 days post inoculation (**Fig. 3A** left panels and **Fig. 4A**, red curves). By contrast, infectious virus titers in female hamsters, who were also challenged with high viral dose, are maximal at inoculation and decay below limit of detection 6-8 days post inoculation (**Fig. 3B** left panels, red curves). The differences in infectious viral dynamics in the female population are due to fewer target cells getting infected compared with males and donors (*T* graphs in **Fig. 5A**, red versus black). Total viral RNA is similar among the high inoculum groups, with peaks trending behind the infectious virus titer peaks by 6 − 12 hours (**Fig. 3** and **Fig. 4** left panel, blue lines).

**Figure 5:**
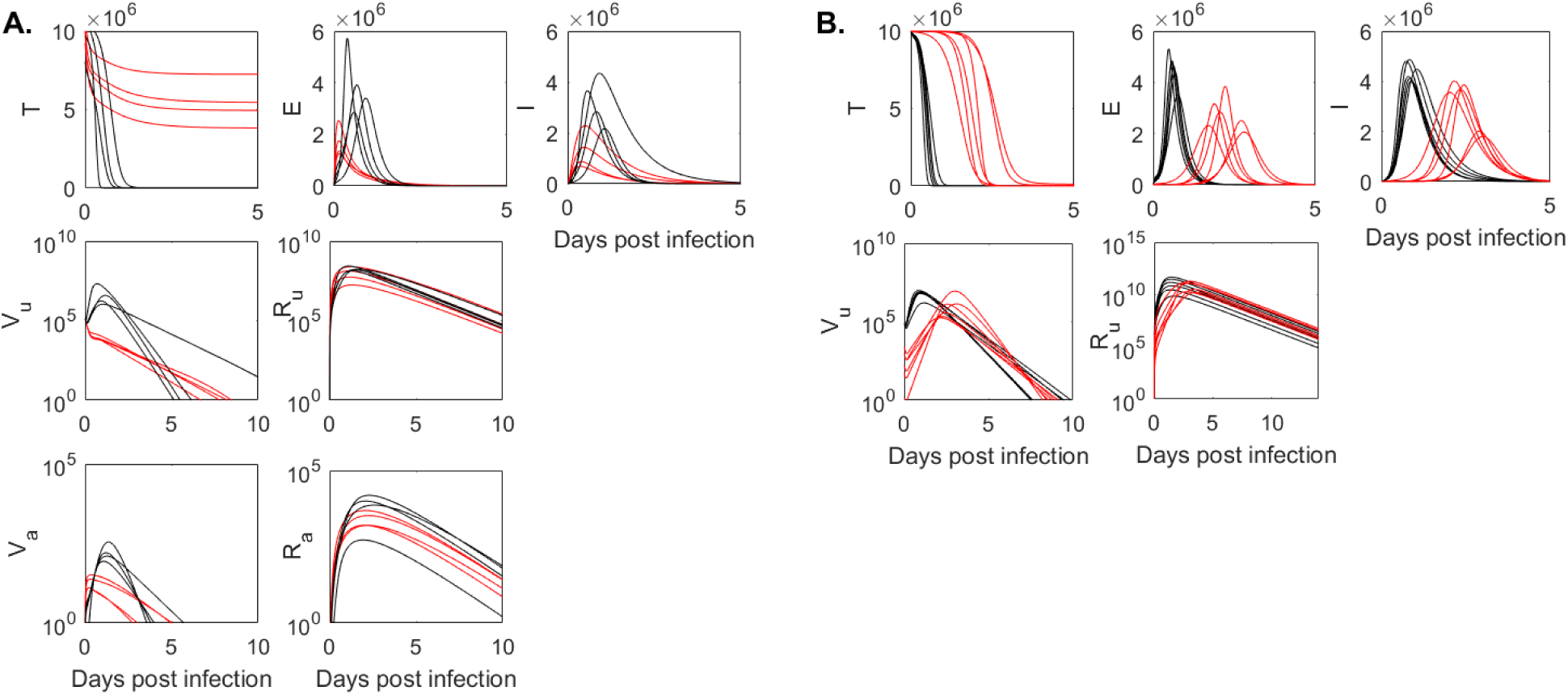
Population dynamics of *T* , *E, I, V*_*u*_, *R*_*u*_, *V*_*a*_ and *R*_*a*_ as given by models Eq. 3 and Eq. 4 in (A.) males (back) and females (red) and (B.) donors (black) and contacts (red). Model parameters are given in Tables 1 and 2.

In the hamsters of the contact group, who were infected (rather than inoculated) with a unknown inoculum dose, infectious virus titers have a delayed expansion compared to the other three groups, peaking 2 − 3 days post infection and decaying below limit of detection 6.7 − 7 days post infection (**Fig. 4B**, red curves). Besides the delay in viral expansion, we observe fewer cells getting infected and a lower overall viremia in contacts compared to donors (**Fig. 5B**, *T* and *V*_*u*_ red versus black graphs). Interestingly, following a delay in expansion, the RNA reach similar levels in contacts and in donors (*R*_*u*_ graphs in **Fig. 5B** red versus black curves).

We find consistent estimates between the groups for the infected cell death rate *δ*, with ranges between 1.78 − 5.44 per day, corresponding to infected cells life-spans of 4.4 − 13.4 hours. The infectivity, *β*, is higher in the female and contact groups, 2.56 × 10^−5^ and 2.07 × 10^−5^ mL/(TCID50 x day), respectively, compared to the male and donor groups, which have similar infectivities of 6.62 × 10^−6^ and 5.37 × 10^−6^ mL/(TCID50 x day), respectively. The production rate of infectious virus is highly variable among groups, with the lowest group average value occurring in females, *p* = 0.1 TCID50/infected cell, followed by contacts, *p* = 9.62 TCID50/infected cell. Males and donors have similar average production rates, 22.3 and 15.6 TCID50/infected cell, respectively. Moreover, the RNA production rate is variable among the two studies but similar among the groups within each study. In particular, *ρ*_1_ = 127 RNA/infected cells and *ρ*_1_ = 214 RNA/infected cells in males and females, and two orders of magnitude higher in donors and contacts, *ρ*_1_ = 8.1×10^4^ RNA/infected cells and *ρ*_1_ = 4.5×10^4^ RNA/infected cells, respectively. The differences between the studies can be noted in the higher total RNA in donors and contacts (**Fig. 4** blue curves), 2.5 order of magnitude hired than in males and females (**Fig. 3** blue curves). Overall infectious virus titers (and consequently theoretical predictions) are two fold higher in donors and contacts compared to males and females, a difference we attribute to the experimental setup in [25] versus [9].

### Viral kinetics in aerosols

In order to determine the relationship between the amount of infectious virus titer (and total viral RNA) shedding into the environment over time and host-viral dynamics in the URT, we added two viral compartments to the within-host model and normalized them to account for subject’s weight variability. The resulting within-host and aerosols model in given by systems **Eq. 3** and **Eq. 4** (see **Materials and Methods** for full derivation). We estimated viral shedding parameters by validating the models against temporal aerosol data (infectious and total RNA) from the males and females (see **Materials and Methods**). The resulting dynamics are in good agreement with the aerosol kinetics in both groups. In particular, the models predict that both infectious virus titers and total viral RNA get released into the air immediately after inoculation. For the males, the shedded infectious virus titers peak 23-29 hours after inoculation, 2-14 hours after the peaks in URT, and decay below limit of detection 2.4-4.9 days after inoculation (**Fig. 3A** right panels, red lines). Total viral RNA peak 1.5-2 days post inoculation, 7-19 hours after the peaks in URT, and persist above limit of detection for the duration of the experiment (**Fig. 3A** right panels, blue lines). For the females, the shedded infectious virus titers are maximal four hours after inoculation and decay below limit of detection 2.3-4 days later, faster than in the male group (**Fig. 3B** right panels, red lines). Females total RNA kinetics, however, are similar to those in male, peaking 18-22 hours after the peaks in URT and persisting above limit of detection for the duration of the experiment (**Fig. 3B** right panels, blue lines). We observe sex-specific differences in the shedding rates, with females infectious virus shedding rate *ϕ*_1_ = 1.9 × 10^−3^ being 5-times higher than that of the males, *ϕ*_1_ = 3.62 × 10^−4^ per day. The RNA shedding rates are similar among the sexes, *ϕ*_2_ = 0.82 × 10^−4^ and 1.02 × 10^−4^ per day for males and females, respectively.

### Basic reproductive ratio

For each group, we estimated the within-host basic reproductive number *R*_0_, which varies over a range between 7.7 and 64 for the male, 0.5 and 0.7 for the females, 16 and 61 for the donors, and 4.4 and 17 for the contacts.

### Relationship between infectious virus in aerosols and transmission

Throughout the course of an infection the hamsters shed both infectious virus and total RNA into the air. Transmission to a close contact occurs when the infectious viruses reach the recipient and establish an infection. We wanted to determine whether aerosol data is a good predictor for the number of infectious virions that jump start such an infection in a close contact. While we know the exact inoculum value for the hamsters in the donor group, we do not know the inoculum value for the hamsters in the contact group. When we include the infectious inoculum *V*_*u*_(0) = *V*_0_ as an unknown parameter to be estimated from the contact hamster data, we find that it varies over a range between 1.4 − 969 TCDI50/ml among the six contacts. Since the contacts were co-housed with the infected donors at day 1 post inoculation, we compared the estimated *V*_0_ with the amount of infectious virus found in the aerosols of males and females. Our model predicted that the aerosol values for the infectious virus titers at day one post inoculation, *V*_*a*_(1), ranged between 96 − 222 TCDI50/ml in males and between 7 − 52 TCDI50/ml in females (see **Fig. 6**). When accounting for the fact that measured infectious virus in the Sia *et al*. study [25] are two-folds higher than the measured infectious virus in the Hawks *et al*. study [9], we find that the aerosol measurements are a good proxy for infectious inoculum.

**Figure 6:**
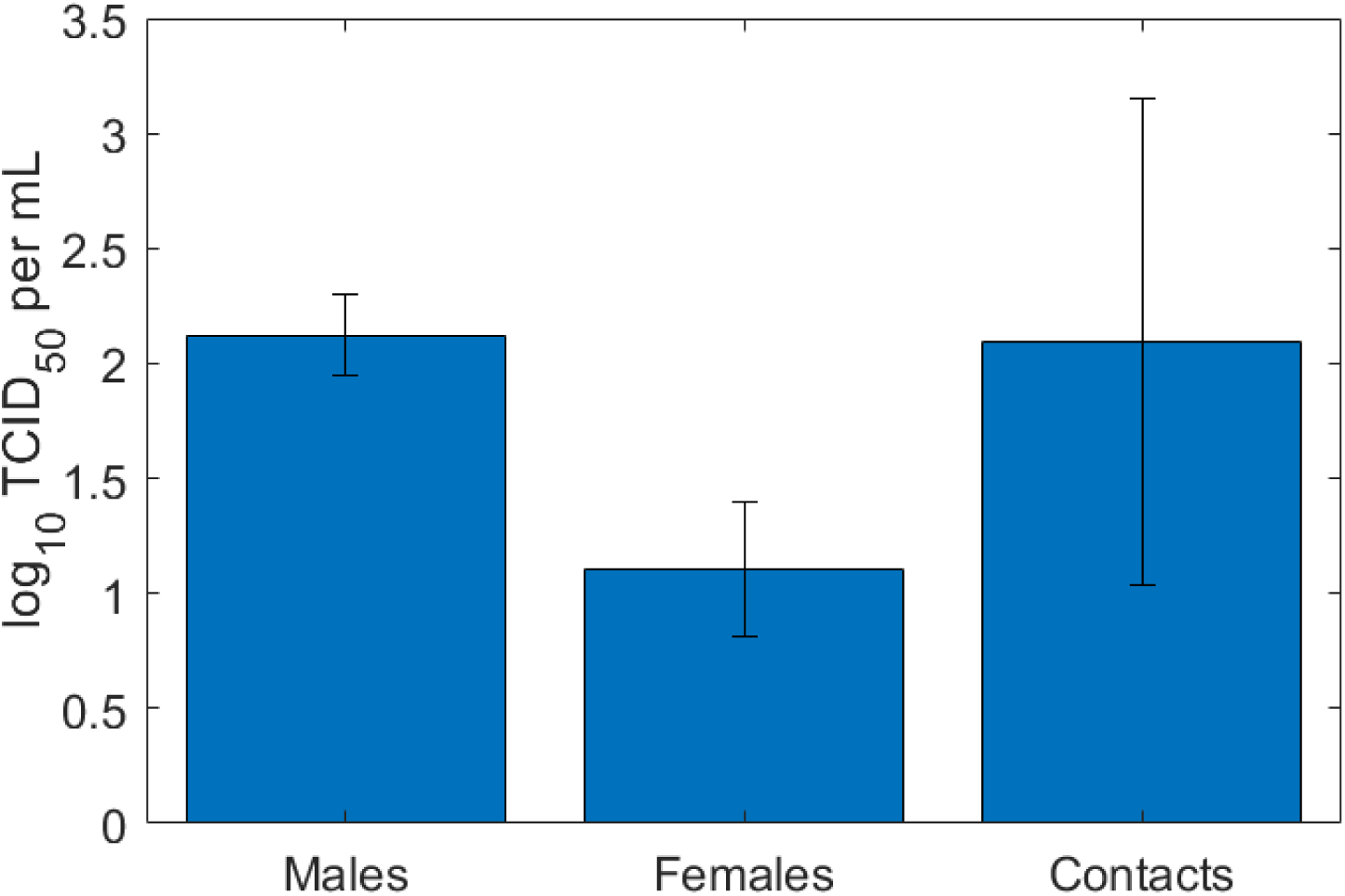
Viral load in aerosols at day one, *V*_*a*_(1), given by models Eq. 3 and Eq. 4 in males and females and *V*_0_ estimates in contacts given by Eq. 3. Model parameters are given in Tables 1 and 2.

**Figure 7:**
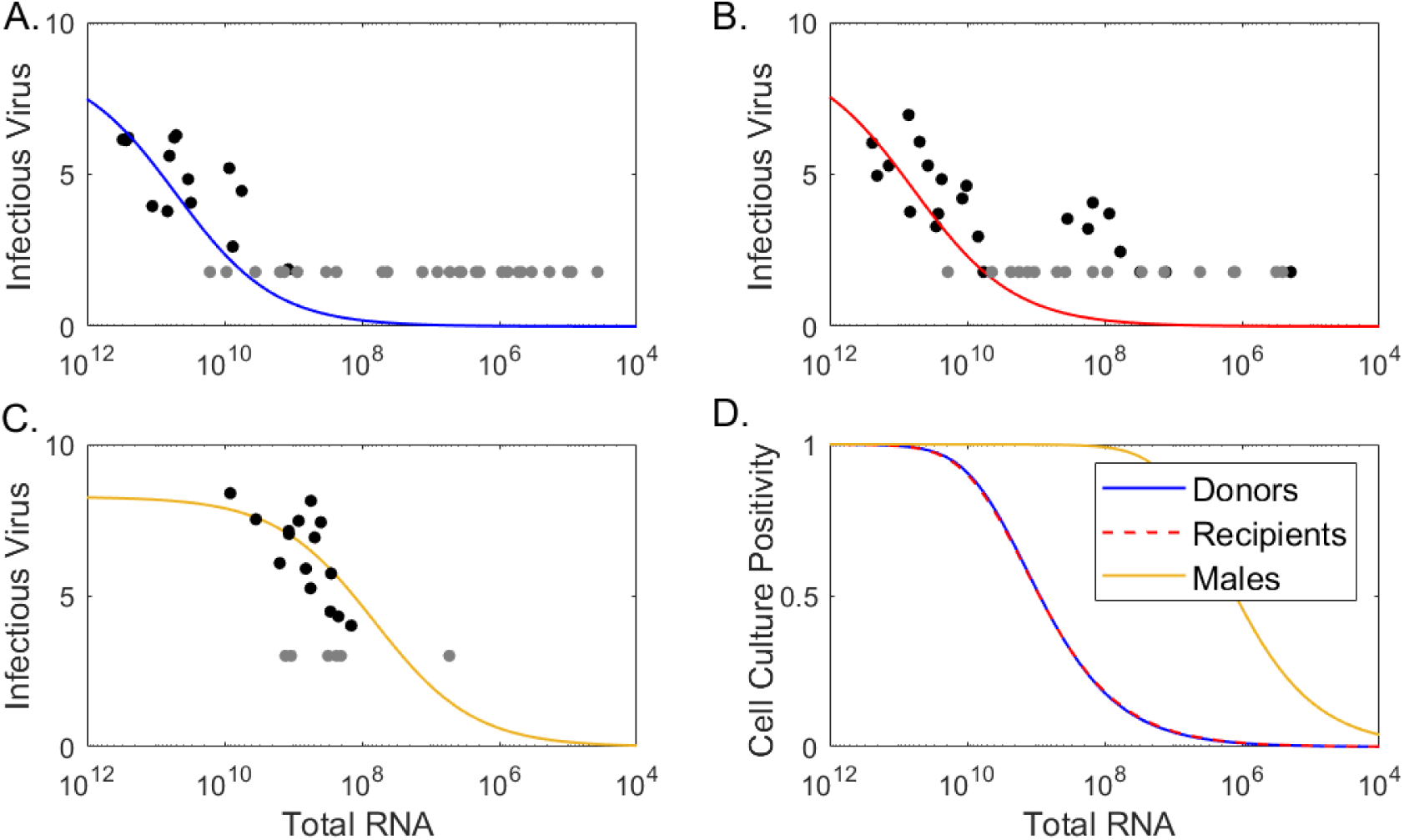
Log10 infectious virus *ν*(*R*_*u*_) as a function of total RNA *R*_*u*_ given by Eq. (7) versus data (circles) in (A.) donors, (B.) contacts, and (C.) males. The grey circles are at the limit of detection and were considered censored data. (D.) Percent cell culture positivity *p*(*R*_*u*_) as a function of total viral RNA *R*_*u*_ given by Eq. (8) for donors (blue line), contacts (red line), and males (gold line). Parameters are given in Table 3.

### Infectious virus versus total RNA levels

Detection of viral RNA by PCR testing from URT samples is the gold standard for COVID-19 diagnosis and is used to instate and discontinue control precautions, such as isolation and quarantine. However, there is no clear correlation between detection of viral RNA and detection of infectious virus titers. We use the data in the two hamster infection studies [9, 25] and our model predictions to investigate: (i) the dependence of infectious viral levels on the RNA levels, and (ii) the dynamics over time of the viral RNA to infectious virus ratio. We exclude the female group from these analyses due to limited amount of infectious virus data above the limit of detection.

To quantify the dependence of infectious viral levels on the viral RNA levels we combined all population (*R*_*u*_, *V*_*u*_) pairs. This resulted in 14 donor, 19 contact, and 15 male pairs above the limit of detection and 24 donor, 17 donor and 6 male pairs below the limit of detection. We assumed that the two quantities can be described by a density dependent function

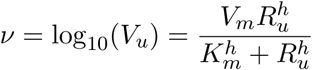

(**Eq. 7** in **Materials and Methods**) as in [8, 13]. Through fitting this function to the population (*R*_*u*_, *V*_*u*_) data in the three groups (taking into consideration the presence of censored data, see **Materials and Methods** for full explanation), we found that the level of infectious viruses increases sub-linearly with increases in viral RNA, with the same exponent h=0.6 in all three groups. The *K*_*m*_ values are three orders of magnitude higher in donors and contacts compared to males, which is due to the higher RNA values in this study.

As in Ke *et al*. [13], we define the probability *p* that a cell culture tests positive to be

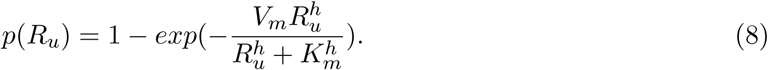

For this assumption and *K*_*m*_ and *h* values estimated from the data, we find an identical probability that a cell culture testing positive in donors and contacts (**Fig. 7D**, red and blue lines) with 10^9^ viral RNA/ml being needed for 50% probability of cell culture positivity. In contrast, 10^6^ viral RNA/ml are enough for 50% probability of cell culture positivity in males (**Fig. 7D**, gold lines).

For each group, we investigated the changes in viral RNA to infectious virus ratio, *R*_*u*_*/V*_*u*_, and found it to be time-dependent, which suggests that the two measurements are explaining different biological processes. Specifically, we found that the average *R*_*u*_*/V*_*u*_ grows between 10^5^ and 10^8^ in the first 4 days in donors, between 2 × 10^5^ and 7 × 10^6^ in the first 5 days in contacts and between 30 and 5 × 10^3^ in the first 3 days in males, consistent with the data (**Fig. 8** blue, red, gold lines). Later *R*_*u*_*/V*_*u*_ ratio predictions from models **Eq**. (3) and **Eq**. (4) are no longer reliable, as the infectious virus decays below limits of detection making the ratio unrealistically high.

**Figure 8:**
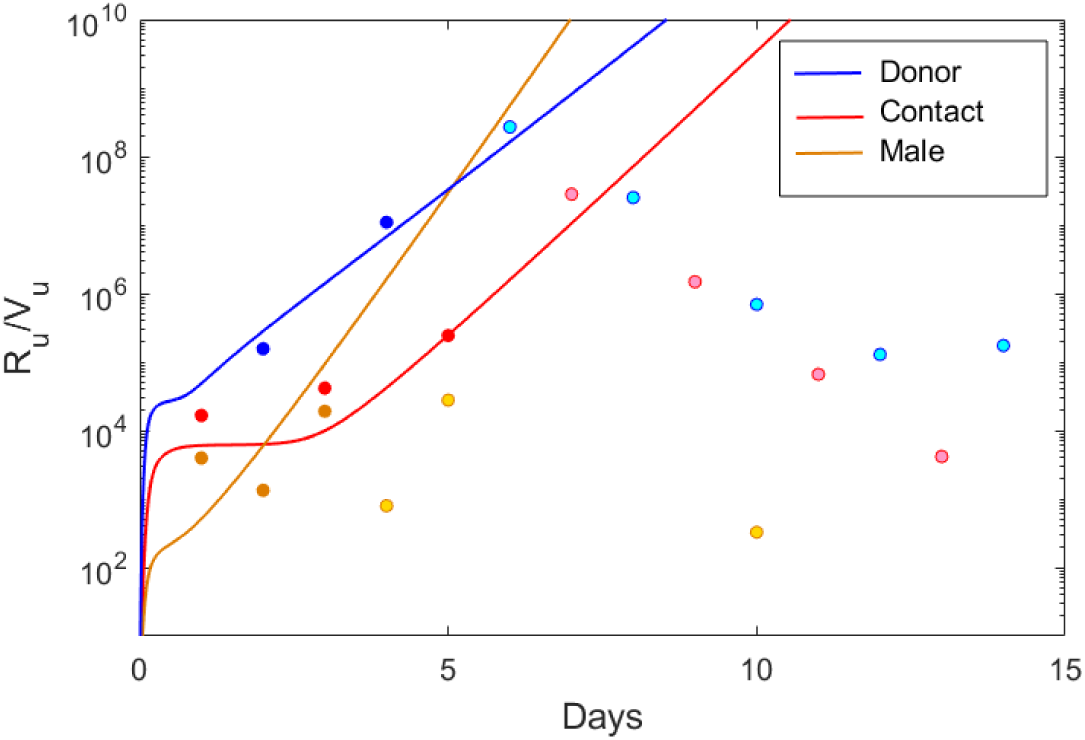
*R*_*u*_*/V*_*u*_ over time versus data in representative donors (blue), contact (red), male (gold) hamsters. The light data points correspond to data where the infectious virus is at the limit of detection.

## 4 Discussion

In this study, we developed within-host and aerosol mathematical models of SARS-CoV-2 infection in golden hamsters that describe the kinetics of infectious virus and viral RNA in the upper respiratory tract and aerosols. We fitted the models to data from two studies [25] and [9] which included four separate groups: three that were challenges with high viral dose (donors, males and females) and one that was challenged with lower viral dose, through infection following close proximity with an infected host (contacts). We estimated several key parameter values for each group and determined inter group variability. We found that the within-host basic reproductive number *R*_0_ is less than one in all female hamsters, indicating limited viral spread. By contrast, *R*_0_ in above one in all male hamsters from all groups (donors, contacts, males), indicating successful viral spread. The hamsters in the contact group had the lowest *R*_0_, ranging between 4.4 − 17, similar to other studies [13]. The other two groups, who were challenged with high viral dose, had larger *R*_0_ estimates, with median *R*_0_ of 31 in donors and 16 in males. The death rate of productively infected cells ranged between 1.78 and 5.44 per day, corresponding to infected cells life-spans of 4.4 to 13 hours, similar within the groups and longer than in humans studies [13].

To model infectious virus shedding into the environment, we extended the upper respiratory tract model to account for infectious virus titer and total RNA emitted into aerosols. Within-host and aerosol model fitting showed that infectious virus titers get released into the air immediately after inoculation with peak shedding 23-29 hours after inoculation in males and 4 hours in females. Infectious viral shedding ends faster in females compared to males, with virus in aerosols losing replication-competency by 2.3-4 days post infection in females and 2.4-4.9 days post inoculation in males. The results are similar with experiments that shown no transmission of SARS-CoV-2 to naive hamsters co-housed with infected donor six days post infection [25].

To determine the relationship between the infectious virus shedding and transmission, we estimated the inoculum dose levels that best describe the kinetic data in the contact group and compared the prediction with the amount of infectious virus present in the aerosol of males and females. When accounting for differences between the studies, we found that aerosol infectious virus titers at day one are a good proxy for the infectious inoculum needed to infect hamsters in the contact group.

Our models and data herein predict that viral RNA persists in both upper respiratory tract and in aerosols long after replication-competent virus stops being detected, with RNA values staying above detectable levels at least a week after the infectious virus is lost. While in public health setting a SARS-CoV-2 diagnostic is determined by PCR assays, RNA levels are not always indicative of virus infectivity, with PCR specificity for detecting replication-competent virus decaying as the cycle threshold (Ct) values increase [4, 4, 18, 32]. We used the models in this study to determine the connection between viral RNA and replication-competent virus levels, and found that the ratio of viral RNA to infectious viral titers is time-dependent and ranges between 10^2^ and 10^8^ RNA/TCID50 in the first five days following infection, wider than in other studies [16,30]. We also found that after day five, the RNA to infectious virus ratio is no longer a reliable measurement of infectiousness, with the measured RNA values indicating the presence of genomic fragments, immune-complexed or neutralised virus, rather than replication-competent virus [2]. These estimates may be species and disease specific, with human studies reporting variable lengths of infectious viral shedding [4,29,31], with larger shedding windows during severe disease [29]. However, our results are consistent with the reported lack of transmission in contact golden hamsters co-housed with an infected donor at day six [25].

We have also investigated the population level relationship between the amount of RNA and the amount of infectious virus in a sample and found that the infectious virus increases in a density-dependent manner with the viral RNA, as suggested in previous work [9,13]. The results (consistent among the three groups considered) showed that when the viral RNA is high, the level of infectious virus saturates, consistent with our temporal results that show that it is unlikely we can predict infectiousness at high RNA:TCID50 ratio. The turning point where viral infectiousness starts to saturate is study dependent, with differences due to experimental settings.

Our study has several limitations. We assumed that both infectious virus clearance and RNA degradation rates are known, with infectious virus clearance set at influenza levels *c* + *d* = 10 per day (corresponding to life-span of 2.4 hours) and the degradation rate set arbitrarily at *d* = 1 per day (corresponding to life-span of one day). Using sensitivity analysis we have found that changing the clearance rates to *c* + *d* = 15 and *c* + *d* = 5 does not influence the results (not shown). In all instances, however, the RNA degradation needs to be small, *d* = 1 or smaller, to explain the differences between infectious virus and viral RNA decay. Moreover, we had too include and estimate an additional removal of the aerosol infectious virus, which we assumed was due to infectious viral inactivation due to the elements. We have also considered that all neutralized virus leads to RNA production. Further information is needed to determine the biological processes leading to increased degradation of infectious virus in aerosols compared to upper respiratory tract and leading to the persistence of RNA in upper respiratory tract and aerosols after infectious virus is lost. Lastly, due to limited aerosol data in the females (with some subjects having just one data point above the limit of detection) we could not properly identify sex-specific differences and excluded this group from some of the analyses.

In conclusion, we have developed a within-host and aerosol model for SARS-CoV-2 infection in golden hamsters and used it to investigate the dynamics of viral RNA and infectious virus titers in URT and aerosols. We validated the models against data and used it to determine the temporal relationship between infectious virus, viral RNA and the probability of a nearby host getting infected. The results can guide interventions.

## Author contributions

**Conceptualization:** Stanca M. Ciupe.

**Formal analysis:** Nora Heitzman-Breen and Stanca M. Ciupe.

**Funding acquisition:** Stanca M. Ciupe.

**Software:** Nora Heitzman-Breen and Stanca M. Ciupe.

**Supervision:** Stanca M. Ciupe.

**Writing – original draft:** Stanca M. Ciupe.

**Writing – review & editing:** Nora Heitzman-Breen and Stanca M. Ciupe.

## Acknowledgements

SMC and SH-B acknowledge funding from National Science Foundation grants No. DMS-1813011 and DMS-2051820 and by a Virginia Tech Center for Emerging, Zoonotic, and Arthropod-borne Pathogens (CeZAP) seed grant. We thank Dr. Nisha Duggal for sharing the data and for helpful discussions on data collection.

## References

[1] Monolix version 2019r2. Antony, France: Lixoft SAS, 2019.

[2] Monique I Andersson, Carolina V Arancibia-Carcamo, Kathryn Auckland, J Kenneth Baillie, Eleanor Barnes, Tom Beneke, Sagida Bibi, Tim Brooks, Miles Carroll, Derrick Crook, et al. Sars-cov-2 rna detected in blood products from patients with covid-19 is not associated with infectious virus. Wellcome open research, 5, 2020.

[3] Martin Z Bazant and John WM Bush. A guideline to limit indoor airborne transmission of covid-19. Proceedings of the National Academy of Sciences, 118(17), 2021.

[4] Jared Bullard, Kerry Dust, Duane Funk, James E Strong, David Alexander, Lauren Garnett, Carl Boodman, Alexander Bello, Adam Hedley, Zachary Schiffman, et al. Predicting infectious severe acute respiratory syndrome coronavirus 2 from diagnostic samples. Clinical infectious diseases, 71(10):2663–2666, 2020.

[5] Jonathan E Forde and Stanca M Ciupe. Modeling the influence of vaccine administration on covid-19 testing strategies. Viruses, 13(12):2546, 2021.

[6] Jonathan E Forde and Stanca M Ciupe. Quantification of the tradeoff between test sensitivity and test frequency in a covid-19 epidemic—a multi-scale modeling approach. Viruses, 13(3):457, 2021.

[7] Antonio Gonçalves, Pauline Maisonnasse, Flora Donati, Mélanie Albert, Sylvie Behillil, Vanessa Contreras, Thibaut Naninck, Romain Marlin, Caroline Solas, Andres Pizzorno, et al. Sars-cov-2 viral dynamics in non-human primates. PLoS computational biology, 17(3):e1008785, 2021.

[8] Ashish Goyal, Daniel B Reeves, E Fabian Cardozo-Ojeda, Joshua T Schiffer, and Bryan T Mayer. Viral load and contact heterogeneity predict sars-cov-2 transmission and superspreading events. Elife, 10:e63537, 2021.

[9] Seth A Hawks, Aaron J Prussin, Sarah C Kuchinsky, Jin Pan, Linsey C Marr, and Nisha K Duggal. Infectious sars-cov-2 is emitted in aerosols. bioRxiv, 2021.

[10] Esteban A Hernandez-Vargas and Jorge X Velasco-Hernandez. In-host mathematical modelling of covid-19 in humans. Annual reviews in control, 50:448–456, 2020.

[11] Adrianne L Jenner, Rosemary A Aogo, Sofia Alfonso, Vivienne Crowe, Xiaoyan Deng, Amanda P Smith, Penelope A Morel, Courtney L Davis, Amber M Smith, and Morgan Craig. Covid-19 virtual patient cohort suggests immune mechanisms driving disease outcomes. PLoS pathogens, 17(7):e1009753, 2021.

[12] Fa-Chun Jiang, Xiao-Lin Jiang, Zhao-Guo Wang, Zhao-Hai Meng, Shou-Feng Shao, Benjamin D Anderson, and Mai-Juan Ma. Detection of severe acute respiratory syndrome coronavirus 2 rna on surfaces in quarantine rooms. Emerging Infectious Diseases, 26(9):2162, 2020.

[13] Ruian Ke, Carolin Zitzmann, David D Ho, Ruy M Ribeiro, and Alan S Perelson. In vivo kinetics of sars-cov-2 infection and its relationship with a person’s infectiousness. Proceedings of the National Academy of Sciences, 118(49), 2021.

[14] Ruian Ke, Carolin Zitzmann, Ruy M Ribeiro, and Alan S Perelson. Kinetics of SARS-CoV-2 infection in the human upper and lower respiratory tracts and their relationship with infectiousness. medRxiv, 2020.

[15] Kwang Su Kim, Keisuke Ejima, Shoya Iwanami, Yasuhisa Fujita, Hirofumi Ohashi, Yoshiki Koizumi, Yusuke Asai, Shinji Nakaoka, Koichi Watashi, Kazuyuki Aihara, et al. A quantitative model used to compare within-host sars-cov-2, mers-cov, and sars-cov dynamics provides insights into the pathogenesis and treatment of sars-cov-2. PLoS biology, 19(3):e3001128, 2021.

[16] William B Klimstra, Natasha L Tilston-Lunel, Sham Nambulli, James Boslett, Cynthia M McMillen, Theron Gilliland, Matthew D Dunn, Chengun Sun, Sarah E Wheeler, Alan Wells, et al. Sars-cov-2 growth, furin-cleavage-site adaptation and neutralization using serum from acutely infected hospitalized covid-19 patients. The Journal of general virology, 101(11):1156, 2020.

[17] YH Li, YZ Fan, L Jiang, and HB Wang. Aerosol and environmental surface monitoring for sars-cov-2 rna in a designated hospital for severe covid-19 patients. Epidemiology & Infection, 148, 2020.

[18] Wang-Da Liu, Sui-Yuan Chang, Jann-Tay Wang, Ming-Jui Tsai, Chien-Ching Hung, Chia-Lin Hsu, and Shan-Chwen Chang. Prolonged virus shedding even after seroconversion in a patient with covid-19. Journal of Infection, 81(2):318–356, 2020.

[19] Eric A Meyerowitz, Aaron Richterman, Rajesh T Gandhi, and Paul E Sax. Transmission of sars-cov-2: a review of viral, host, and environmental factors. Annals of internal medicine, 174(1):69–79, 2021.

[20] Shelly L Miller, William W Nazaroff, Jose L Jimenez, Atze Boerstra, Giorgio Buonanno, Stephanie J Dancer, Jarek Kurnitski, Linsey C Marr, Lidia Morawska, and Catherine Noakes. Transmission of sars-cov-2 by inhalation of respiratory aerosol in the skagit valley chorale superspreading event. Indoor air, 31(2):314–323, 2021.

[21] Teresa Moreno, Rosa María Pintó, Albert Bosch, Natalia Moreno, Andrés Alastuey, María Cruz Minguillón, Eduard Anfruns-Estrada, Susana Guix, Cristina Fuentes, Giorgio Buonanno, et al. Tracing surface and airborne sars-cov-2 rna inside public buses and subway trains. Environment international, 147:106326, 2021.

[22] Jin Pan, Seth A Hawks, Aaron J Prussin, Nisha K Duggal, and Linsey C Marr. Sars-cov-2 on surfaces and hvac filters in dormitory rooms. Environmental Science & Technology Letters, 2021.

[23] Mehrshad Sadria and Anita T Layton. Modeling within-host sars-cov-2 infection dynamics and potential treatments. Viruses, 13(6):1141, 2021.

[24] Ye Shen, Changwei Li, Hongjun Dong, Zhen Wang, Leonardo Martinez, Zhou Sun, Andreas Handel, Zhiping Chen, Enfu Chen, Mark H Ebell, et al. Community outbreak investigation of sars-cov-2 transmission among bus riders in eastern china. JAMA internal medicine, 180(12):1665–1671, 2020.

[25] Sin Fun Sia, Li-Meng Yan, Alex WH Chin, Kevin Fung, Ka-Tim Choy, Alvina YL Wong, Prathanporn Kaewpreedee, Ranawaka APM Perera, Leo LM Poon, John M Nicholls, et al. Pathogenesis and transmission of sars-cov-2 in golden hamsters. Nature, 583(7818):834–838, 2020.

[26] Amber M Smith, Frederick R Adler, Julie L McAuley, Ryan N Gutenkunst, Ruy M Ribeiro, Jonathan A McCullers, and Alan S Perelson. Effect of 1918 pb1-f2 expression on influenza a virus infection kinetics. PLoS computational biology, 7(2):e1001081, 2011.

[27] Naveen K Vaidya, Angelica Bloomquist, and Alan S Perelson. Modeling within-host dynamics of sars-cov-2 infection: A case study in ferrets. Viruses, 13(8):1635, 2021.

[28] Neeltje Van Doremalen, Trenton Bushmaker, Dylan H Morris, Myndi G Holbrook, Amandine Gamble, Brandi N Williamson, Azaibi Tamin, Jennifer L Harcourt, Natalie J Thornburg, Susan I Gerber, et al. Aerosol and surface stability of sars-cov-2 as compared with sars-cov-1. New England journal of medicine, 382(16):1564–1567, 2020.

[29] Jeroen JA van Kampen, David AMC van de Vijver, Pieter LA Fraaij, Bart L Haagmans, Mart M Lamers, Nisreen Okba, Johannes PC van den Akker, Henrik Endeman, Diederik AMPJ Gommers, Jan J Cornelissen, et al. Duration and key determinants of infectious virus shedding in hospitalized patients with coronavirus disease-2019 (covid-19). Nature communications, 12(1):1–6, 2021.

[30] Elisa Vicenzi, Filippo Canducci, Debora Pinna, Nicasio Mancini, Silvia Carletti, Adriano Lazzarin, Claudio Bordignon, Guido Poli, and Massimo Clementi. Coronaviridae and sars-associated coronavirus strain hsr1. Emerging infectious diseases, 10(3):413, 2004.

[31] Sunpeng Wang, Yang Pan, Quanyi Wang, Hongyu Miao, Ashley N Brown, and Libin Rong. Modeling the viral dynamics of sars-cov-2 infection. Mathematical biosciences, 328:108438, 2020.

[32] Roman Wölfel, Victor M Corman, Wolfgang Guggemos, Michael Seilmaier, Sabine Zange, Marcel A Müller, Daniela Niemeyer, Terry C Jones, Patrick Vollmar, Camilla Rothe, et al. Virological assessment of hospitalized patients with covid-2019. Nature, 581(7809):465–469, 2020.

[33] Lian Zhou, Maosheng Yao, Xiang Zhang, Bicheng Hu, Xinyue Li, Haoxuan Chen, Lu Zhang, Yun Liu, Meng Du, Bochao Sun, et al. Breath-, air-and surface-borne sars-cov-2 in hospitals. Journal of aerosol science, 152:105693, 2021.

